# Mesenchymal Cell-Derived Extracellular Vesicles Ameliorate Age-Related Deficits in Working Memory as well as Brain MRI and CSF *in vivo* Biomarkers of Neurodegeneration in Rhesus Monkeys

**DOI:** 10.1101/2024.11.14.623673

**Authors:** Evan C. Mackie, Chia-Hsin Cheng, Maya Alibrio, Christine Rutledge, Hongqi Xin, Michael Chopp, Ryan McCann, Douglas L. Rosene, Qiong Yang, Ella Zeldich, Maria Medalla, Bang-Bon Koo, Tara L. Moore

**Affiliations:** Department of Anatomy & Neurobiology, Boston University Chobanian & Avedisian School of Medicine, 72 E Concord St, Room L1004, Boston, MA USA 02118; Center for Systems Neuroscience, Boston University, 610 Commonwealth Ave, 7th Floor, Boston, MA USA 02215; Department of Neurology, Henry Ford Health, 2799 W Grand Blvd, Detroit, MI USA 48202; Graduate Program in Neuroscience, Boston University, 610 Commonwealth Ave, Boston, MA USA 02118; Department of Biostatistics, Boston University School of Public Health, 715 Albany St, Boston, MA USA 02118

## Abstract

Normal aging in humans and non-human primates is associated with a decline in cognitive functions. Subject-wise differences in cognitive decline can be attributed to different degrees of damage to cortical white matter (WM) which is largely affected by neuroinflammation during aging. Mesenchymal stromal cell-derived extracellular vesicles (MSC-EVs) have recently been identified as a potential immunomodulatory therapeutic for brain damage and Alzheimer’s disease (AD) and related dementias by suppressing neuroinflammation. Here, we evaluated the efficacy of MSC-EVs for slowing or ameliorating cognitive decline during aging in rhesus monkeys, a well-studied model of normal aging that is free of extensive AD pathology. We report that late middle-aged monkeys treated with MSC-EVs every two weeks for 18 months showed improved performance on a task of spatial working memory relative to vehicle control monkeys. In addition, we used diffusion magnetic resonance imaging (MRI) and resting state functional MRI to evaluate structural white matter and functional network changes in vivo. Imaging data revealed that MSC-EV treatment preserved prefrontal and temporal WM structural integrity and large-scale functional network connectivity that are correlated with early, increased CSF levels of amyloid beta protein. Amyloid beta levels at 12 months are also correlated with improved cognitive performance at the end of the 18 months of treatment. These findings suggest that MSC-EVs can mitigate age-related cognitive decline by potentially enhancing the CSF clearance of neurodegenerative proteins, which correlates with greater WM integrity and functional brain connectivity.

## 1. Introduction

Normal aging is characterized by deficits in the cognitive domains of learning and memory and executive function (Albert et al., 2007; Bachevalier, 1993; Bartus et al., 1978, 1979; Blacker et al., 2007; Henry et al., 2004; Light, 1991; Moore et al., 2003, 2006; Presty et al., 1987; Steere and Arnsten, 1997). These changes are not a consequence of widespread cortical neuronal loss, but rather are thought to result from myelin pathology, synapse loss and inflammation predominantly in the frontal and medial temporal lobes (Bowley et al., 2010; Brickman et al., 2011; DeVries et al., 2024; Moore et al., 2005; Peters and Sethares, 2002; Peters et al., 2001; Shobin et al., 2017; Wisco et al., 2008). While numerous studies have investigated age-related neuroinflammation, decreases in gray and white matter volume, and degenerative changes in these regions (Arnsten et al., 1994; Arnsten and Goldman-Rakic, 1990; Arnsten and Jentsch, 1997; Grady, 1998; Makris et al., 2007; Moore et al., 2005; Peters et al., 1994; Raz et al., 1997; Sawaguchi et al., 1990; reviewed in West, 2000; West, 1996), fewer studies have focused on the occurrence of and relationship between amyloid beta (Aβ) protein deposition and changes in white matter integrity and their possible role underlying age-related cognitive decline. Increasingly, there is evidence of Aβ protein accumulation and deposits in cognitively-intact aged individuals, which has been described as a pathological change in normal aging (Jonkman et al., 2020; reviewed in Mormino and Papp, 2018). Rodrigue et al., 2012, demonstrated that in a large sample of cognitively intact adults, aged 60 and over, there was markedly elevated Aβ deposition that was related to subtle cognitive changes. Other studies have revealed that many cognitively normal elderly evidence Aβ neuropathology upon autopsy with Aβ deposition being most prominent in the frontal, cingulate, and medial temporal regions (Crystal et al., 1993; Hulette et al., 1998; Jack et al., 2008; Knopman et al., 2003; Morris and Price 2001; Price and Morris 1999). While Aβ deposition appears to occur with normal aging, the exact extent remains unknown, and whether it plays a role in normal cognitive aging remains unclear.

Alterations in white matter are well documented in the normal-aged human and monkey brain. A review by Bennett and Madden, 2014, provided extensive evidence in humans that cerebral white matter integrity declines in healthy aging, as measured by diffusion tensor imaging, that are likely the result of changes in underlying myelin. This is supported by a study with 100 individuals in a European consortium project that showed a relationship between age, cognitive performance, and a degradation in white matter integrity (Coelho et al., 2021). Further, in aged rhesus monkeys, myelin integrity in the prefrontal cortex and frontal white matter tracts declines, as shown by electron microscopy and reduced fractional anisotropy with diffusion magnetic resonance imaging (MRI) (Bowley et al., 2010; Makris et al., 2007; reviewed in Freire-Cobo et al., 2021; Luebke et al., 2010). Additionally, white matter volume decreases with age, with the most pronounced loss occurring in the frontal lobe (Kubicki et al., 2019; Wisco et al., 2008; reviewed in Luebke et al., 2010). In conjunction with this white matter pathology in normal aging, age-related depositions of neurogenerative proteins, such as Aβ, also occur in humans and monkeys (Gearing et al., 1996; Jonkman et al., 2020; reviewed in Stonebarger et al., 2021). While white matter pathology has been shown to be strongly correlated with cognitive decline (Bowley et al., 2010; Peters et al., 1994; Peters et al., 2000; Peters et al., 2001), it remains unclear how levels of Aβ reflect the severity of cognitive impairments associated with age and neurodegenerative disease. Thus, there is a critical need to understand how these age-related pathological changes interact and drive cognitive aging in order to develop effective therapeutics to reverse these changes.

One potential therapeutic is mesenchymal stromal/stem cell-derived extracellular vesicles (MSC-EVs) (Golpanian et al., 2017; Larrick and Mendelsohn, 2017; Sanz-Ros et al., 2022b; Schulman et al., 2018; Tompkins et al., 2017; reviewed in Heldring et al., 2015; Sanz-Ros et al., 2022a). Extracellular vesicles (EVs) released by MSCs carry bioactive proteins, microRNAs, and lipids and show promise as a therapeutic for reversing age-related changes in the brain (see reviews by Heldring et al., 2015; Kou et al., 2022). MSC-EVs have demonstrated these effects in a wide variety of animal models of disease and injury, especially for reducing inflammation, neurodegeneration, and facilitating repair. Our group has previously shown the efficacy of MSC-EVs in a rhesus monkey model of cortical injury in ameliorating injury-related microglial inflammatory phenotypes, enhancing neuronal and myelin plasticity, and diminishing oxidative damage (Go et al., 2020; Go et al., 2021; Medalla et al., 2020; Zhou et al., 2023). MSC-EVs have also been shown to be effective in mitigating chronic neurodegeneration. For example, in rodent models of Alzheimer’s disease (AD), there is evidence of MSC-EVs restoring cognitive function, enhancing neurogenesis, decreasing pro-inflammatory cytokines (e.g. TNF-α, IL-1β, and IL-6) and microglia activation, increasing anti-inflammatory cytokines (e.g. IL-10, IL-4, and IL-13), and reducing Aβ concentrations and plaque deposition (Cui et al., 2019; Ding et al., 2018; Li, B. et al., 2020; Losurdo et al., 2020; Ma et al., 2020; Reza-Zaldivar et al., 2019; Wang and Yang, 2021; Yang et al., 2020; reviewed in Chakari-Khiavi et al., 2019; Gonçalves et al., 2023; Guo et al., 2020; Liew et al., 2017; Reza-Zaldivar et al., 2018; Sanz-Ros et al., 2022a). Additionally, evidence in the literature has shown a potential protective effect of stem cell derived EVs, as they contain antioxidants and Aβ degrading enzymes such as neprilysin and insulin degrading enzyme (Apodaca et al., 2021; Bodart-Santos et al., 2019; de Godoy et al., 2018; Ding et al., 2018; Elia et al., 2019; Lee et al., 2018; reviewed in Sanz-Ros et al., 2022a). Our group has also shown that MSC-EV-treatment can mitigate AD-related cellular phenotypes and decrease pathological Aβ and p-tau depositions in an *in vitro* 3D cortical spheroid (CS) model generated from Down Syndrome patient-derived induced Pluripotent Stem Cells (iPSCs) (Campbell et al., 2023).

Together, the evidence from our laboratory and from the literature overwhelmingly support the premise that MSC-EVs play a role in mitigating inflammation and promoting clearance of damaged proteins that likely facilitate myelin repair and reduce pathology in *in vitro* and *in vivo* animal models of disease and injury. However, few studies have directly evaluated the ability of MSC-EVs to alleviate age-related cognitive decline and the associated pathological changes in the aging brain, particularly in primate models. To begin to fill this gap in our knowledge, in the current study, we investigated how chronic, systemic administration of MSC-EVs affects age-related cognitive decline and underlying changes in neuropathology and structural connectivity in our rhesus monkey model of aging. Our data provide evidence that in late middle-aged monkeys, longitudinal systemic treatment with MSC-EVs (every two weeks for 18 months) can reverse or slow age-related cognitive decline while increasing levels of age-associated neurodegenerative proteins released into the cerebrospinal fluid (CSF) and strengthening white matter integrity and functional connectivity.

## 2. Methods

### 2.1. Subjects

Six late middle-aged rhesus monkeys (ages 17-24 years of age, equivalent to 51-72 human years) were used in this study (Table 1). All monkeys had known birth dates, complete health records, and were obtained from National Primate Research Centers or private vendors. All monkeys received medical examinations before entering the study. In addition, explicit criteria were used to exclude monkeys with a history of splenectomy, thymectomy, exposure to radiation, organ transplantation, malnutrition, chronic illness including viral or parasitic infections, neurological diseases or chronic drug administration. Each of the monkeys underwent an initial structural MRI to ensure there was no occult neurological damage. Results of the evaluations revealed that all monkeys were healthy at the time they were entered into the study.

**TABLE 1.** Biographical details of subjects used in this study.

| SUBJECT | SEX | GROUP | DOB | BASELINE TESTING | RETESTING | AGE AT BASELINE TESTING | AGE AT RETESTING |
| --- | --- | --- | --- | --- | --- | --- | --- |
| AM380e | Male | Vehicle | 5/22/96 | 10/1/20 | 1/30/23 | 24.4 | 26.7 |
| SM071 | Female | Vehicle | 2/1/03 | 12/1/20 | 4/20/23 | 17.8 | 20.2 |
| AM384e | Male | Vehicle | 11/10/03 | 1/19/22 | 2/1/24 | 18.2 | 20.2 |
| AM381e | Male | MSC-EV | 3/1/96 | 10/1/20 | N/A | 24.6 | N/A |
| AM383e | Female | MSC-EV | 3/1/00 | 3/1/21 | 4/4/23 | 21.0 | 23.1 |
| AM382e | Female | MSC-EV | 6/3/00 | 6/1/21 | 9/28/23 | 21.0 | 23.3 |

(N/A = Subject AM381e was removed before the end of the study due to declining health. DOB = Date of Birth)

While on study, monkeys were individually housed in colony rooms in the Boston University Animal Science Center where they were in constant auditory and visual range of other monkeys. This facility is fully AAALAC accredited, and animal maintenance and research were conducted in accordance with the guidelines of the National Institutes of Health and the Institute of Laboratory Animal Resources Guide for the Care and Use of Laboratory Animals. All procedures were approved by the Boston University Institutional Animal Care and Use Committee. Diet consisted of Lab Diet Monkey Chow (#5038—Lab-Diet Inc., St. Louis, MO) supplemented by fruit and vegetables with feeding taking place once per day, immediately following cognitive testing. During testing, small pieces of fruit or candy were used as rewards. Water was available continuously in the monkeys’ home cages, and monkeys were housed under a 12-hour light/dark cycle with cycle changes occurring in a graded fashion over the course of an hour. Monkeys were checked daily by trained observers for health and well-being and were given a medical exam every three months by a clinical veterinarian.

### 2.2. General Study Design

All six monkeys received baseline MRI, CSF collections, and cognitive testing consisting of initial Delayed Non-Matching to Sample (DNMS) acquisition, 5 days each of DNMS 2- and 10-minute delays, and 10 days of testing on the Delayed Recognition Span Task – Spatial (DRSTsp). Monkeys were then randomly assigned to the treatment (2 females and 1 male) and vehicle (1 female and 2 males) groups and began receiving intravenous (IV) infusions of either MSC-EVs or vehicle once every two weeks for 18 months (Table 1). Each monkey in the treatment group received a dose of 4 × 10^11^ MSC-EVs at each infusion, while monkeys in the vehicle group received an IV infusion of vehicle phosphate-buffered saline (PBS) that did not contain MSC-EVs. During the treatment period, CSF was collected every three months and MRIs were repeated every 6 months. At the completion of the 18 months of treatment, monkeys were re-tested on cognitive tasks while continuing to receive infusions of MSC-EVs or vehicle every two weeks. All study personnel were blind to group assignments throughout the study.

Due to limited availability of animals, we were unable to balance the groups for sex. Future studies will involve adding additional animals to fully assess sex differences in treatment efficacy.

### 2.3. Baseline Cognitive Testing

Prior to beginning treatments (control (vehicle) or MSC-EV administration), all monkeys were initially familiarized with cognitive testing in a Wisconsin General Testing Apparatus (WGTA) where they were trained to displace a single gray plaque to obtain a reward. On each trial, the plaque was placed pseudo-randomly over one of the three food wells, each 3.5 cm in diameter and 0.5 cm deep. Wells were spaced 11.5 cm apart in a single row across the middle of a testing board (See Figure 1A). Small pieces of fruit or candy were used as rewards during testing. Monkeys were trained until they responded on 40 consecutive trials on each of two successive days. Then monkeys completed the DNMS (acquisition and 2- and 10-minute delays) and the DRSTsp.

**Figure 1.**
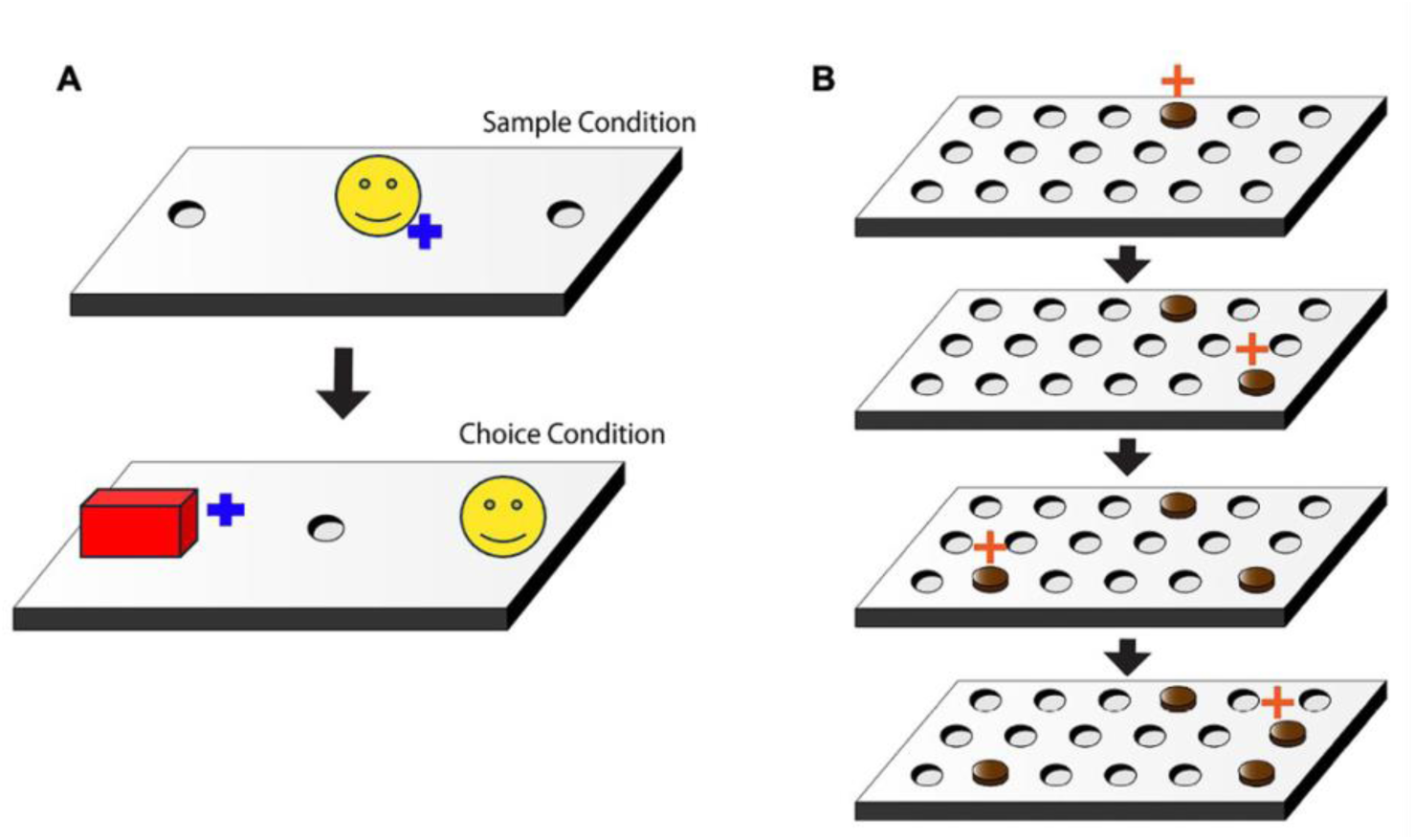
Schematic of the Delayed Non-Matching to Sample and Delayed Recognition Span Task - Spatial. **A)** A schematic showing the sample and choice presentation during one trial of the Delayed Non-Matching to Sample (DNMS) task. In the sample condition, the monkey must displace the object presented over the center well to retrieve a food reward (blue cross). After a delay (10, 120, or 600 seconds), the monkey must now choose the new, unfamiliar object placed over a pseudo-randomly chosen lateral well to obtain the food reward (blue cross). **B)** A schematic showing sequential stimuli (brown disc) presentation within one trial of the Spatial Delayed Recognition Span Task (DRSTsp). During each presentation the monkey must choose the disc in the new spatial location (red cross). Each successive correct response trial was followed by the addition of a new disc in a novel location on the testing board and this continued until the monkey made an error (i.e., chose a previously chosen disc). With the occurrence of the first error, the trial was terminated and the number of discs on the testing board minus one were counted to determine the recognition span score for that trial (i.e., number of correct consecutive responses).

### 2.4. Initial Acquisition – Delayed Non-Matching to Sample (DNMS)

The DNMS task is a benchmark recognition memory task that assesses the ability of subjects to identify a novel stimulus from a familiar stimulus following a 10-sec delay interval. In the present study, all monkeys began the acquisition phase of the DNMS test in the WGTA. Using the same testing board as familiarization testing (Figure 1A), each trial began with a sample object presented over the central baited food well. The monkey was permitted to displace the object and obtain the reward. The WGTA door was then lowered, and the original, now familiar, sample object was placed over an unbaited lateral well and a novel, unfamiliar object was placed over the other lateral well that was baited. Ten seconds after the original sample trial, the choice trial was begun, and the monkey was required to choose the unfamiliar, novel object to obtain the reward. Both testing objects were replaced after each trial so as not to reuse objects during the same testing period. Ten seconds later, the next trial was initiated with a new, novel sample object presented over the baited central well. Ten seconds later, another recognition trial was given using the second sample object and another new novel object. The position of the two objects on successive recognition trials was varied from left to right lateral wells in a pre-determined pseudorandom order. A non-correction procedure was used and twenty trials per day were given until the monkey reached a learning criterion of 90 correct responses in 100 consecutive trials or a maximum of 2000 trials. Objects were drawn from a pool of 600 “junk” objects and paired so that in each daily session of 20 trials, 60 of the objects were used. Once all the initial pairings were used, (30 days of 20 trials per day), the 600 objects were randomly recombined to produce new pairs so that the pairings presented continued to be new and unique on each trial.

### 2.5. Delayed Non-Matching to Sample (DNMS) 2-Minute Delays

Upon reaching criterion on the basic task, the 10 sec delay between the presentation of the sample object and the recognition trial was increased to 120 seconds. Ten trials a day for 5 days (for a total of 50 trials) at the delay interval were administered with the monkey remaining in the testing apparatus during the delay interval.

### 2.6. Delayed Non-Matching to Sample (DNMS) 10-Minute Delays

Upon completion of the trials with the 2-minute delay between the presentation of the sample object and the recognition trial, the delay was increased to 600 seconds. Five trials a day for 5 days (for a total of 25 trials) at the delay interval were administered with the monkey remaining in the testing apparatus during the delay interval.

### 2.7. Delayed Recognition Span Task – Spatial (DRSTsp)

The testing board for the DRSTsp was similar to the DNMS board (Figure 1), but instead of a single row of three wells, this board had three rows of six wells each (3.5cm wide, 0.5cm deep) and the wells were spaced 6cm apart within a row. The rows were spaced 1.5cm apart. For this task, fifteen identical plain brown discs (6 cm in diameter) were used as stimuli. During the first sequence of a trial, one disc was placed over one of the eighteen wells, which was baited with a food reward. The WGTA door was then raised, and the monkey was allowed to displace the disc to obtain the reward. The door was then lowered, the first disc was returned to its original position over the now unbaited well and a second disc was placed on the board over a baited well in a different location. After 10 seconds, the door was once again raised, and the monkey was required to identify the new second disc in its novel spatial location to obtain the reward. Each successive correct response trial was followed by the addition of a new disc in a novel location on the testing board and this continued until the monkey made an error (i.e. chose a previously chosen disc). With the occurrence of the first error, the trial was terminated and the number of discs on the testing board minus one were counted to determine the recognition span score for that trial (i.e. number of correct consecutive responses). Five such trials were presented each day for ten consecutive days (total 50 trials).

### 2.8. MRI Scanning

Each monkey underwent baseline structural MRI to ensure there was no occult neurological damage, and scans were repeated every 6 months during treatment until the end of the 18-month period of the study. All scans were conducted on the 3 Tesla whole body wide bore Philips Ingenia Elition scanner (Center for Biomedical Imaging, Boston University School of Medicine). Due to a scanner upgrade that occurred mid-study, we have focused on analyzing MRI data collected at the end of the 18-month treatment duration to ensure consistency in data.

For each scan session, monkeys were placed into a specially designed acrylic stereotaxic device that was created to allow for the imaging of monkeys using the 8-channel, phase-array Philips head coil. This ensures homogeneity of the MR field and even application of the signal from the radio frequency pulse (i.e. no drop off in signal) along with access to the standard set of filters and image acquisition tools on the scanner. For scans, monkeys were initially sedated with Ketamine (10mg/kg IM) that was followed with additional doses, as needed.

The following scans were acquired in each MRI session: 1) A structural MRI was acquired using a T1-weighted 3D-turbo field echo (TFE) sequence that was fully optimized to provide high signal to noise ratio (SNR) and high gray-white matter contrast at high resolution with following parameters: Repetition time (TR)/Echo time (TE) = 7 ms/3 ms, flip angle = 8°, number of excitations/averages (NEX) = 8, voxel size: 0.7×0.7×0.7 mm^3^, sagittal plane acquisition. 2) Diffusion MRI (dMRI) was collected using a diffusion weighted imaging spin-echo (DWIse) sequence modified based on the Adolescent Brain Cognitive Development (ABCD) multi-band (MB) diffusion sequence with the following parameters: TR/TE = 4200ms/67ms, flip angle = 90°, number of directions = 108, b-values = 500, 1000, 2000, 3000m/s, voxel size = 1.875×1.875×2mm^3^. 3) Resting state functional MRI (rs-fMRI) was collected from a Fast Field Echo (FFE) single-shot echo planar imaging (EPI) sequence with the following parameters: TR/TE = 2000ms/30ms, flip angle = 52°, slice thickness=2.34mm, 30 slices, field of view (FOV) = 150×150×70.19mm^3^, and 200 volumes.

### 2.9. Cerebrospinal Fluid (CSF) Collection

CSF collected from each monkey at baseline, every 3 months during MSC-EV or vehicle administration, and at euthanasia. For CSF collection, monkeys were sedated with Ketamine (10mg/kg) and the head stabilized in a stereotactic head holder that allows the head to be rotated ventrally relative to the spinal cord. The atlanto-occipital junction was palpated, the head rotated forward to enlarge the target, and the cisterna magna was gently penetrated with a 23G needle and 2mL of CSF was collected. CSF was centrifuged to precipitate contaminants and aliquoted into cryotubes and frozen and stored at −80°C until used for ELISA analysis.

### 2.10. Mesenchymal Stromal Cell-Derived Extracellular Vesicle (EV) Preparation and Administration

EVs were isolated from bone marrow MSCs of a young monkey, as described previously (Moore et al., 2019; Xin et al., 2013). Briefly, a young monkey (approximately 6 years old - equivalent to 18 human years) was sedated with Ketamine (10 mg/kg IM), then anesthetized with sodium pentobarbital (15–25 mg/kg IV). Bone marrow was extracted from the iliac crest and shipped on wet ice with same day delivery to Henry Ford Health in Detroit. Upon arrival, marrow was spun at 4000 x g for 15 min to separate cells. The buffy coat was discarded, and remaining cells were washed in culture medium. Cells were then plated in a T75 flask using media containing 20% FBS and alpha-MEM, grown to confluence, and passaged as necessary. To grow enough cells for EVs, 10 × 10^6^ cells were seeded into a Quantum Incubator (Terumo BCT, Lakewood, CO) and grown in alpha-MEM with 10% EV-depleted FBS (Systems Biosciences, Palo Alto, CA). To harvest EVs, media was collected every other day for four days, then every day for two days. As described in Zhang et al., 2015, media was centrifuged in multiple stages at 250 x g for 5 min, then 3000 x g for 30 min, then filtered and centrifuged a final time at 100,000 x g for 2 h to pellet EVs. The pellet was resuspended in a small volume of PBS. EV concentrations and particle sizes were measured using a qNano (Izon, Cambridge, MA), then diluted to 4 × 10^11^ particles for each monkey in 10 mL of PBS. EVs had an average diameter of 111 nm in size, and had confirmed expression of CD9, CD81, and CD63 by Western blot, per MISEV guidelines (data not shown) (Witwer et al., 2013). The dose was chosen based on previous dosing studies conducted in rats (Xin et al., 2013; Zhang et al., 2015). Following resuspension, MSC-EV or PBS vehicle vials were individually labeled with subject identifiers, stored at −80°C, and shipped to Boston University. The day prior to intravenous administration in monkeys, treatment vials were allowed to thaw overnight at 4°C.

### 2.11. MSC-EV Administration

Following the completion of baseline testing, the monkeys were randomly assigned to either the MSC-EV treatment or control (vehicle) groups and began receiving IV dosing of MSC-EVs or vehicle once every two weeks. For each administration, monkeys were sedated with Ketamine (10mg/kg, IM) and then an IV catheter was inserted into a branch of the saphenous vein. An infusion set was then used to administer 10ml of MSC-EV (4×10^11^ particles in 10mL of PBS) or vehicle. Dosing was continued for 18 months.

At the end of the 18-month dosing period, the monkeys began re-testing on the DNMS task and the DRSTsp while continuing to receive the biweekly doses of MSC-EVs or vehicle (control). Re-acquisition of the DNMS task was administered as described above and monkeys were re-tested until reaching a criterion level of 90 correct responses in 100 consecutive trials. The DNMS delays and DRSTsp were administered as described above.

### 2.12. ELISA CSF Biomarker Analysis

CSF aliquots were thawed just before analysis and all ELISAs were performed according to manufacturer’s instructions with minor optimization of sample dilutions. Samples were analyzed using the Invitrogen Human Aβ40 (KHB3481), Invitrogen Human Aβ42 (KHB3441), Uman Diagnostics Neurofilament light (NFL) (10-7002), and the Ansh Labs Myelin Basic Protein (MBP) (AL-108) assay kits. All plates were read on a Spectramax M3 (Molecular Devices) microplate reader at wavelengths specified by the manufacturers.

### 2.13. MRI Processing and Analysis

Prior studies from our group have utilized a combination of various diffusion and functional MRI techniques and advanced reconstruction methods to assess white matter (WM) integrity, functional network connectivity, and structural-functional properties in the aging rhesus monkey brain. In an earlier study focused on middle aged monkeys, we observed weaker structural connectivity between the hippocampus and the frontal cortex, as examined with diffusion spectrum imaging, accompanied by stronger functional connection strengths between the regions of interests (ROIs) as examined by independent component analysis which were shown to correlate with worse performance on the DRSTsp (Koo et al., 2013). In the current study, we sought to expand on these findings by adopting a model-free diffusion reconstruction method, q-space sampling (Yeh et al., 2010), to detect more detailed microstructural white matter changes. Recent studies with resting-state fMRI have defined large-scale functional brain networks (e.g. default mode network, executive network, salience network, etc.) in the human brain by grouping brain regions that share similar functional properties into the same network and have a lesser focus on anatomical organization and boundaries (Yeo et al., 2014) which have been modified onto the rhesus monkey brains (Xu et al., 2019). In this study, we focused on changes in WM integrity, resting state networks (RSN), and the relationships between MRI biomarkers, cognitive performance outcomes, and CSF biomarkers to better understand the therapeutic role of MSC-EVs.

All raw MRI images were converted and preprocessed using a modified in-house pipeline (Koo et al., 2013). In this study, T1 and rs-fMRI data were processed using the AFNI (analysis of functional NeuroImages) macaque fMRI processing pipeline (Jung et al., 2021), which provided hierarchical cortical regions of interest (ROI) parcellation that can be further mapped into major resting state functional networks (RSNs) by matching anatomical and neurological proximity between NHP and human brain functional networks (Xu et al., 2019). 5 RSNs were extracted from the rs-fMRI and 10 inter-network connectivity (i.e., connection between RSNs) were calculated from the pipeline. A total of 15 functional network features were used for the analysis. Diffusion MRI processing included generalized q-space imaging (GQI) reconstruction from DSI Studio, a model-free method to evaluate microstructural diffusivity measured as the generalized fractional anisotropy (GFA) (Yeh et al., 2010). B0 images were then registered to the standard NIMH Macaque Template (NMT) brain template. A diffusion-tensor imaging (DTI)-based white matter (WM) atlas containing 76 WM tracts were then inverse transformed to the dMRI space to provide matching anatomical definitions in the reconstructed GFA maps (Zakszewski et al., 2014).

### 2.14. Statistical Analysis

Analyses of group differences in cognitive testing scores were conducted using one-way analyses of variance (ANOVA) in GraphPad Prism (version 10.2.3 for macOS, GraphPad Software, Boston, Massachusetts USA). Analyses of longitudinal CSF protein concentrations were conducted using two-way repeated measures ANOVA (group x timepoint) with Tukey’s multiple comparisons or paired t-test for post-hoc analyses. Linear correlations between CSF and cognitive data were conducted using Pearson correlations in GraphPad Prism (10.2.3 for macOS). MRI group differences between EV-treated and control subjects were performed with two-sample t-test using MATLAB (R2024a, The MathWorks Inc. Natick, MA). Pairwise linear correlation analyses were performed between 1) multimodal MRI measures, 2) MRI and cognitive testing measures, and 3) MRI and CSF biomarker measures. Pearson correlation coefficient (r) or t-value is reported along with p-value for significant comparisons (MATLAB, R2024a, The MathWorks Inc. Natick, MA). All statistical tests were performed with an alpha level of 0.05 for statistical significance and adjusted for age and sex.

## 3. Results

### 3.1. Baseline Cognitive Testing

Prior to beginning intravenous administration of MSC-EV or vehicle, monkeys completed baseline testing on the DNMS task (basic and delays) and the DRSTsp. There was no significant group difference on any of the tasks completed during the baseline testing after adjusting for age and sex.

### 3.2. Re-testing on Cognitive Battery

Following 18 months of biweekly administration of MSC-EVs or vehicle, when monkeys were not tested, repeat testing began on the DNMS task (basic and delays) and the DRSTsp. All monkeys successfully reacquired criterion (90% correct over 100 trials) on the DNMS basic task, then completed DNMS with 2- and 10-minute delays, and finally the DRSTsp task.

### 3.3. Delayed Non-Matching to Sample (DNMS) Re-acquisition

The total number of trials and errors to criterion were analyzed for each monkey for the reacquisition of the DNMS basic task (Table 2). Separate one-way repeated measures ANOVAs were conducted for trials and for errors with treatment group as the between subject variable. There was no significant overall effect of treatment group for total number of trials to criterion [F (1, 4) = 0.610, p = 0.478] and no significant overall effect of group for total errors to criterion [F (1, 4) = 1.12, p = 0.349].

**TABLE 2.** Subject cognitive performance data.

| SUBJECT | SEX | GROUP | DRSTsp TOTAL SPAN | DRSTsp PEs | DRSTsp PEs DIFFERENCE FROM BASELINE | DRSTsp TOTAL SPAN DIFFERENCE FROM BASELINE |
| --- | --- | --- | --- | --- | --- | --- |
| AM380e | Male | Vehicle | 1.76 | 27 | -22.73 | -2.67 |
| SM071 | Female | Vehicle | 2.2 | 29 | -38.10 | -2.83 |
| AM384e | Male | Vehicle | 2.24 | 29 | -7.41 | 0.33 |
| AM381e | Male | MSC-EV | 2.64 | 10 | 68.75 | 3.83 |
| AM383e | Female | MSC-EV | 2.64 | 18 | 37.93 | 2.67 |
| AM382e | Female | MSC-EV | 2.38 | 16 | 23.81 | 4.00 |

(DRSTsp = Spatial Delayed Recognition Span Task. PEs = Perseverative Errors)

A difference score between performance at baseline and the retesting after treatment was also calculated for each monkey to determine whether there were differences in the change of performance on the DNMS task from baseline to re-testing (Table 2). Separate one-way ANOVAs revealed that there was no significant group difference in the change of performance from baseline to the re-testing for trials [F (1, 5) = 1.71, p = 0.261] or errors [F (1, 5) = 0.993, p = 0.375] on DNMS. These results suggest that there was no effect of MSC-EVs on performance on the DNMS task.

### 3.4. Delayed Non-Matching to Sample (DNMS) Delays

The percent of correct responses on the DNMS basic task with 2-minute and 10-minute delays was determined for each monkey (Table 2). Separate one-way ANOVAs with group as the between subject variable revealed no significant effect of group at 2-minute [F (1, 4) = 0.151, p = 0.718] or 10-minute delays [F (1, 4) = 0.500, p = 0.519].

A difference score was also calculated for the performance of each monkey to determine their change in performance between baseline and the retesting (Table 2). Separate one-way ANOVAs with group as the between subject variable revealed no significant effect of group on the change of performance from baseline to re-testing for 2-minute [F (1, 4) = 0.682, p = 0.455] and 10-minute delays [F (1, 4) = 0.0438, p = 0.845]. These results, taken together with the results from the DNMS basic task, suggest that systemic infusions of MSC-EVs biweekly over an 18-month period does not have a significant effect on recognition memory following either a relatively short (10 sec) or longer (120 or 600 sec) delay.

### 3.5. Delayed Recognition Span Task – Spatial (DRSTsp)

The mean total span achieved by each monkey was determined across the ten days on the DRSTsp at re-testing (Table 2). A one-way ANOVA with treatment group as the between subject variable revealed a significant overall effect of treatment group on DRSTsp performance [F (1, 4) = 7.60, p = 0.05] as shown in Figure 2A. This analysis was followed by a one-way ANOVA to determine if significant group differences in change of performance from baseline to re-testing were evident. As shown in Figure 2C, this analysis revealed a significant group difference on performance change between baseline and re-testing on the DRSTsp [F (1, 4) = 22.098, p = 0.0093] with treated monkeys performing at a higher level on the DRSTsp at re-testing relative to their baseline performance. In contrast, the performance on the DRSTsp at retesting by the monkeys that received vehicle only was either similar to their performance at baseline or had decreased in terms of spatial memory span. These results suggest that the monkeys that received MSC-EVs demonstrated a greater improvement in performance from baseline to re-testing than the control (vehicle) monkeys on the DRSTsp, a task of working memory.

**Figure 2.**
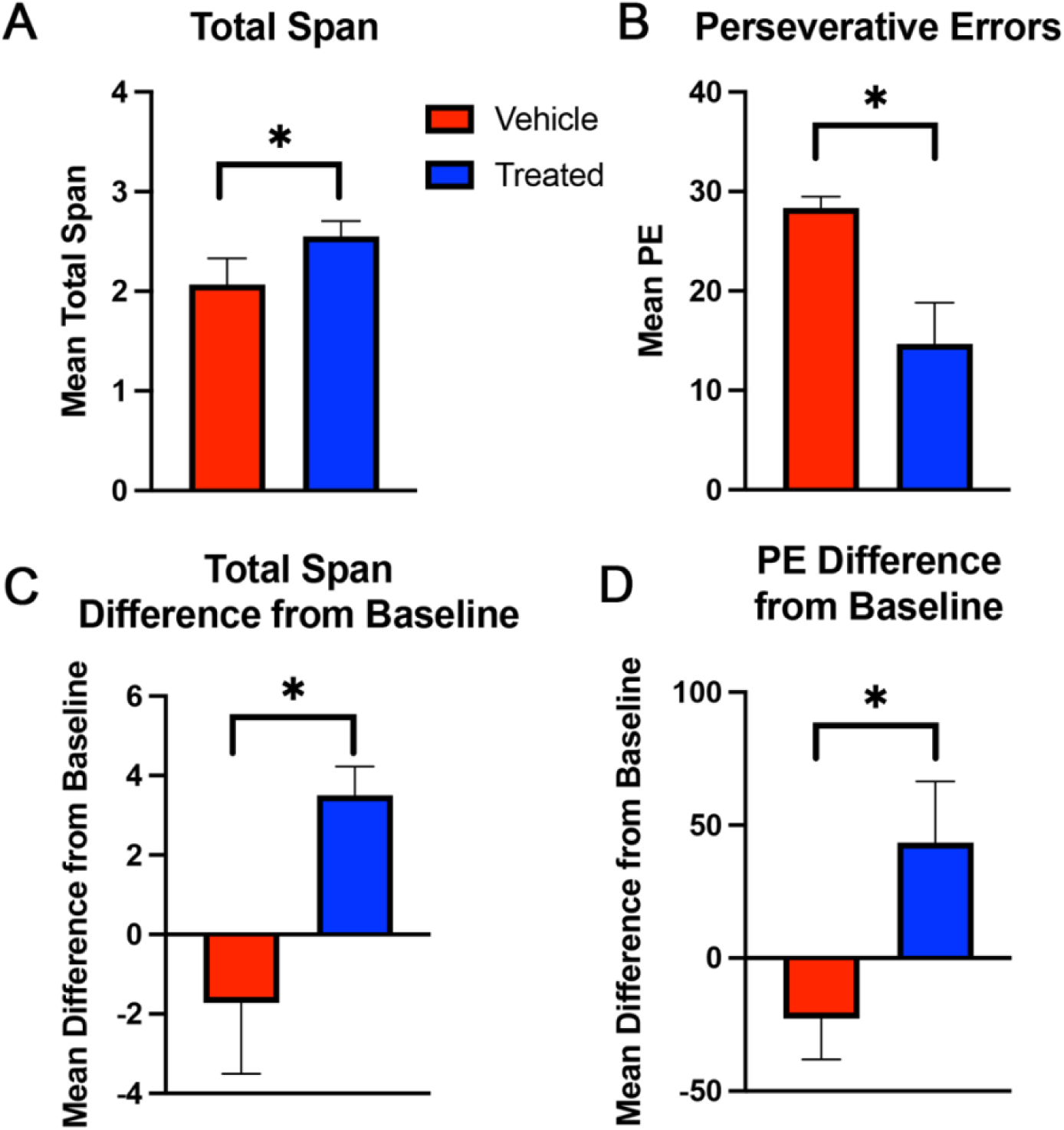
Cognitive Performance on the Delayed Recognition Span Task - Spatial. **A)** MSC-EV treated monkeys achieved a significantly higher span score on the DRSTsp at retesting as compared to vehicle treated monkeys [F (1, 4) = 7.60, p = 0.05]. **B)** MSC-EV treatment significantly reduced the frequency of perseverative responses on the DRSTsp at retesting [F (1, 4) = 30.018, p = 0.005]. **C)** When comparing span scores between baseline and retesting, MSC-EV treated monkeys had significantly higher spans on the DRSTsp after treatment [F (1, 4) = 22.098, p = 0.0093]. **D)** When comparing the frequency of perseverative responses between baseline and retesting, MSC-EV treated monkeys demonstrated significantly fewer preservative errors after treatment [F (1, 4) = 17.239, p = 0.01].

Finally, during testing on the DRSTsp, the number of perseverative errors (when the monkey made an error by choosing the previously correct disc) was determined. A one-way ANOVA revealed a significant effect of MSC-EVs on the number of perseverative errors [F (1, 4) = 30.018, p = 0.005] (Figure 2B) with treated monkeys making fewer perseverative errors than monkeys that received vehicle only. There was a significant group effect in the difference in the number of perseverative errors between baseline and re-testing [F (1, 4) = 17.239, p = 0.01] (Figure 2D). Treated monkeys made significantly fewer perseverative errors at re-testing than at baseline, while monkeys that received only the vehicle made more perseverative errors at re-testing than at baseline. This finding suggests that monkeys that received MSC-EVs for 18 months demonstrated less perseverative response tendencies than monkeys that received vehicle.

### 3.6. CSF Protein Biomarkers of Neurodegeneration

To understand the temporal dynamics of MSC-EV treatment, we monitored protein biomarkers at various time points during treatment. Protein concentrations of biomarkers of neurodegeneration were assessed in CSF collected prior to the start of treatment and then every three months during treatment for 18 months. Levels of AD-related amyloid beta isoforms, Aβ40, Aβ42, and markers of myelin and axonal degeneration, MBP, and NFL, were quantified and compared between groups across the 18-month period using ELISAs (Figures 3 and 4). No significant differences were found between groups using two-way repeated measures ANOVA in any of the measured absolute analyte concentrations at the timepoints analyzed (Figure 3). However, when quantifying protein levels as a percent change from baseline (Figure 4), MSC-EV treated monkeys had significantly higher levels of Aβ42 in CSF after 12 months of treatment as compared to control monkeys (t-test, p = 0.0474) (Figure 4B). Further, linear regression and correlation analysis showed that both Aβ40 (r = 0.861, p = 0.028) and Aβ42 (r = 0.95, p = 0.004) levels at 12 months were positively correlated with changes between baseline and terminal DRSTsp performance (Figure 5). These findings suggest that MSC-EV treatment may enhance cognitive performance through increasing Aβ levels in the CSF.

**Figure 3.**
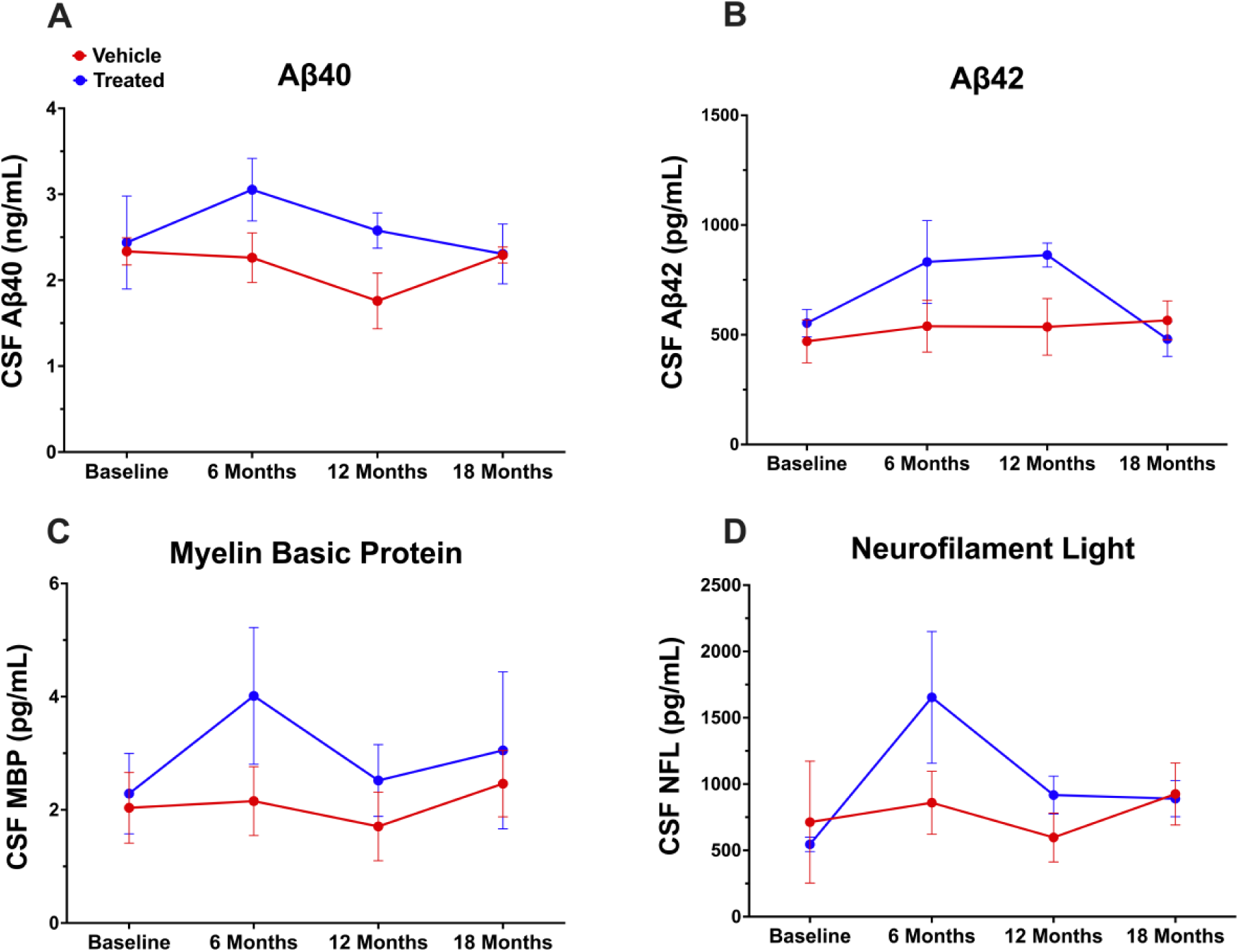
CSF protein markers of neurodegeneration over the course of MSC-EV treatment. Mean protein concentrations of **A)** Aβ40, **B)** Aβ42, **C)** MBP, and **D)** NFL were determined in longitudinal CSF samples with ELISA. No significant group differences were observed at any timepoint.

**Figure 4.**
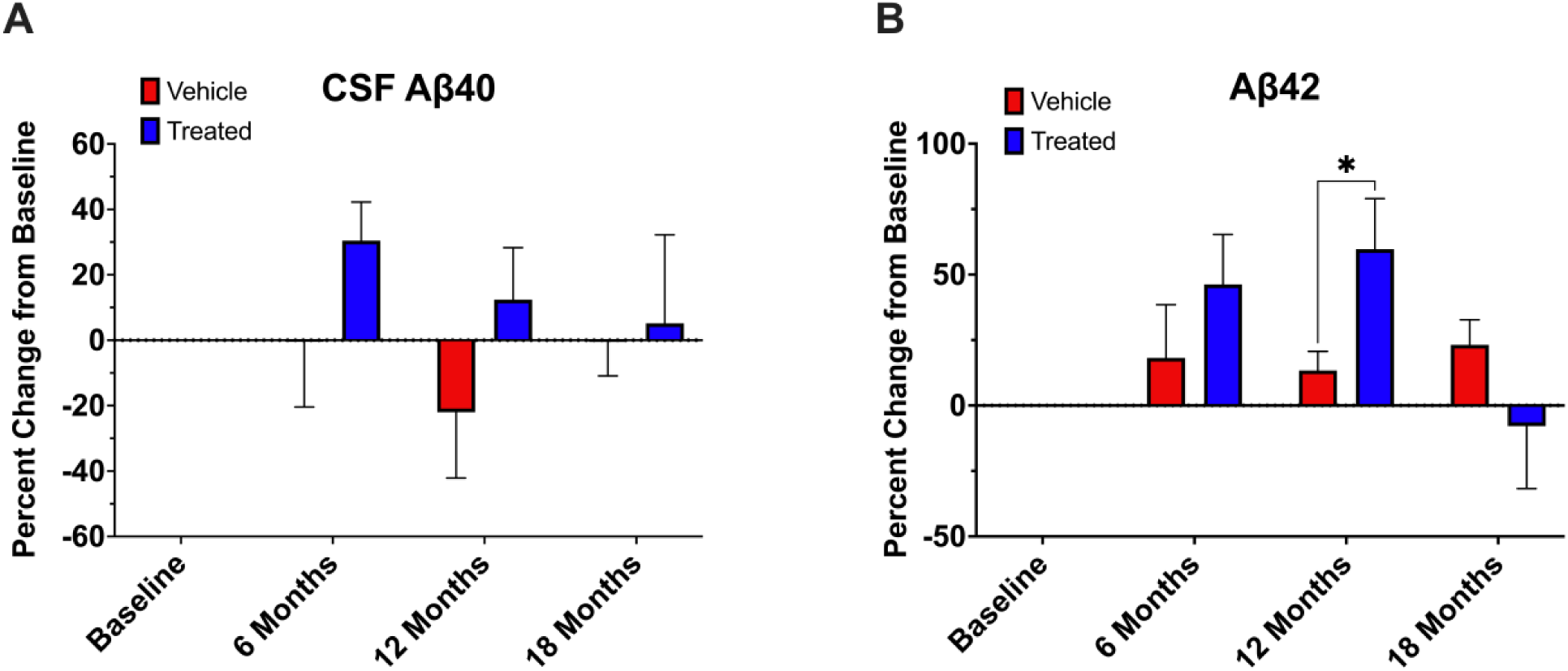
Percent change in CSF concentrations of Aβ as compared to baseline. **A)** Percent change in mean Aβ40 concentration from baseline – no significant differences were observed. **B)** Percent change in mean Aβ42 concentration from baseline – MSC-EV treated monkeys had significantly higher change in CSF Aβ42 protein as compared to vehicle treated monkeys at 12 months of treatment (Student’s t test, p = 0.0474).

**Figure 5.**
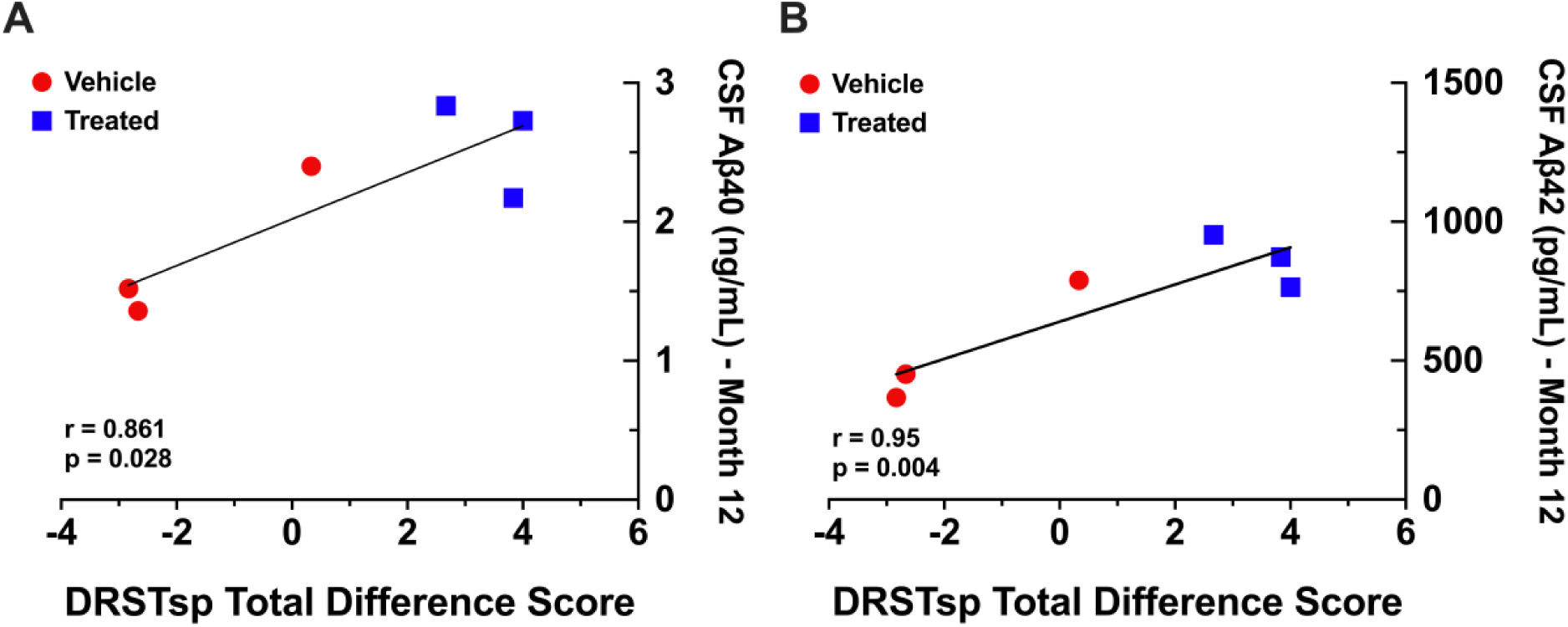
Correlations between DRSTsp performance and CSF Aβ levels. DRSTsp difference scores between baseline and retesting are significantly positively correlated with **A)** CSF Aβ40 (r = 0.861, p = 0.028) and **B)** Aβ42 (r = 0.95, p = 0.004) levels at 12 months after beginning MSC-EV treatment.

### 3.7. MRI Markers of Brain Structural and Functional Properties

To compare brain structural and functional network changes *in vivo* between groups, we used diffusion MRI and rs-fMRI data acquired after 18 months of treatment. Diffusion WM integrity was analyzed via GFA mapping, which was significantly higher in the EV-treated group compared to the control group in the left tapetum (Student’s t test, p = 0.04), the left superior temporal gyrus (Student’s t test, p = 0.029), and the right middle temporal gyrus (Student’s t test, p = 0.009) (Fig. 6A). Resting state network (RSN) connectivity analysis on rs-fMRI data revealed significantly weaker connections in the EV-treated monkeys between the limbic to executive network (Student’s t test, p = 0.029) and between the default mode to salience network (Student’s t test, p = 0.014) (Fig. 6B). WM GFA in the right middle temporal gyrus is negatively correlated with connectivity strength between the default mode and salience network (r = −0.89, p = 0.017), with EV-treated monkeys showing higher WM GFA and lower between-network connectivity (Fig. 6C).

**Figure 6.**
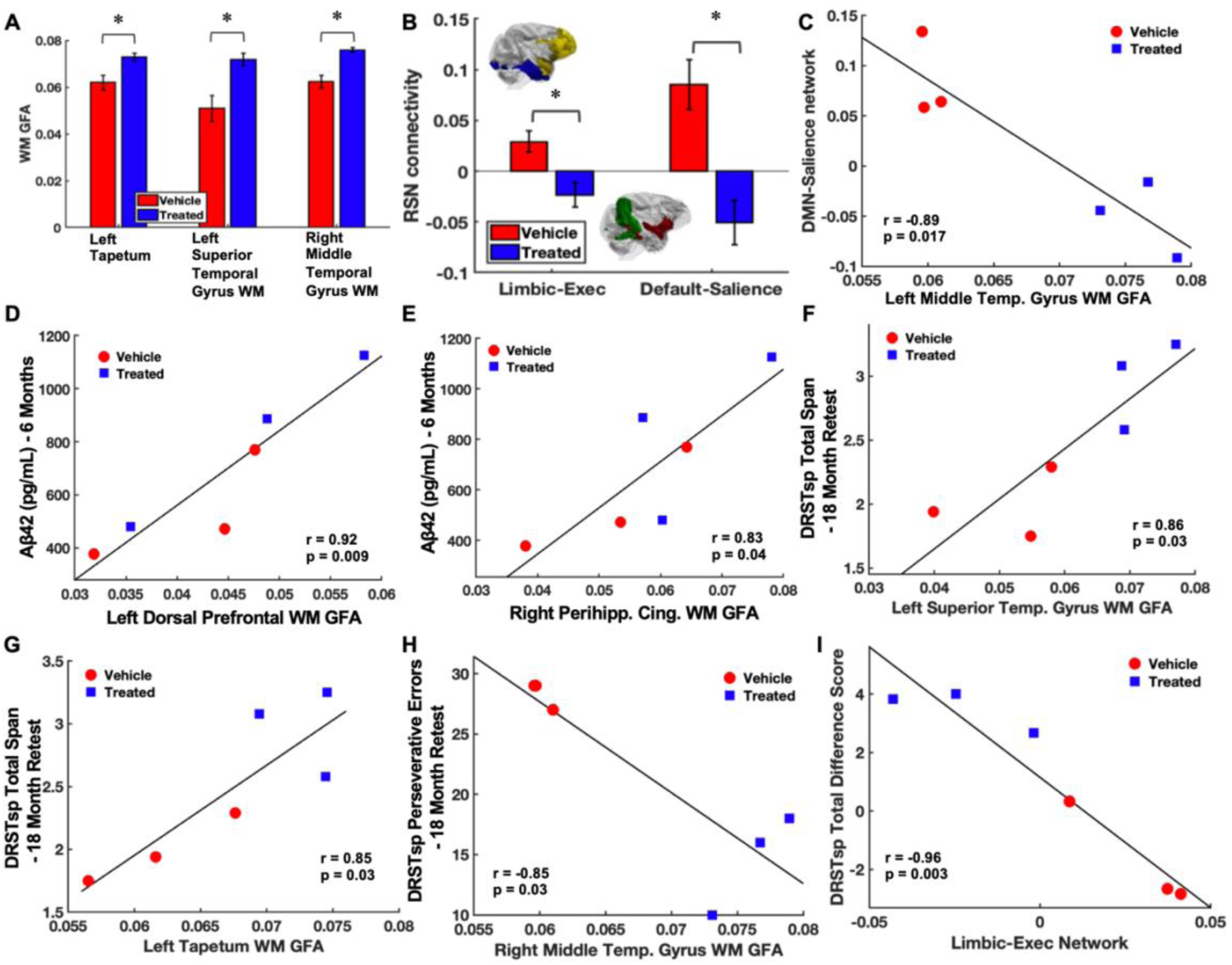
MRI outcomes at the end of 18-month MSC-EV treatment. **A)** Temporal brain regions showed group differences in white matter (WM) generalized fractional anisotropy (GFA) in the MSC-EV treated group, indicating better WM integrity. **B)** Functional MRI shows hyper-connectivity between the limbic and executive networks (Limbic-Exec), and default mode and salience networks (Default-Salience) in the non-treated group. **C**) Decreased DMN-Salience network functional connectivity in MSC-EV treated monkeys correlates with better temporal WM structural integrity. **D&E**) Enhanced CSF Aβ42 clearance at 6 month correlates with better diffusion WM integrity shown as higher GFA in the frontal and temporal WM regions. **F&G**) Increased WM integrity in the temporal WM regions were correlated with better performance on the DRSTsp span task. **H**) Increased WM integrity in the temporal WM regions were correlated with less preservation errors in the DRSTsp task. **I**) Decreased limbic-executive network functional connectivity in MSC-EV treated monkeys correlated with an improved DRSTsp Span task difference score from baseline to terminal testing

We then compared MRI measures of brain structural and functional network integrity to CSF biomarkers of neurodegeneration and cognitive performance. Correlation analysis demonstrated positive relationships between CSF Aβ42 levels at 6 months and diffusion GFA measured in the left dorsal prefrontal (r = 0.92, p = 0.009) WM and right perihippocampal cingulum bundle (r = 0.83, p = 0.04) (Fig. 6D&E). In addition, positive correlations were found between 18-month DRSTsp re-test scores and WM GFA in the left tapetum (r = 0.85, p = 0.03) and the left superior temporal gyrus (r = 0.86, p = 0.03) (Fig. 6F&G), as well as a negative correlation between the 18-month DRSTsp preservative errors and WM GFA in the right middle temporal gyrus (r = −0.85, p = 0.03) (Fig 6H). DRSTsp difference scores were also negatively associated with RSN connectivity between the limbic to executive network (r = −0.96, p =0.003) (6I). Together, MRI findings showed more well-preserved structural WM and resting state functional network integrity in EV-treated animals, which were correlated with Aβ clearance into the CSF and cognitive performance, respectively.

## 4. Discussion

### 4.1. Summary of Results

The current study utilized cognitive testing, diffusion and functional MRI analyses, and longitudinal assessment of protein biomarkers of neurodegeneration in CSF to investigate the effects of MSC-EV treatment on age-related changes in cognition, neuropathology, and white matter integrity and brain functional network connectivity in late middle-aged rhesus monkeys. The overall findings of this study were that long-term MSC-EV treatment: 1) Improves overall spatial working memory performance; 2) Reduces the frequency of perseverative responses on a spatial working memory task; 3) Increases Aβ42 protein concentration, possibly due to enhanced clearance, in the CSF which is correlated with DRSTsp performance; and 4) Preserves prefrontal and temporal WM structural integrity and large-scale functional network connectivity that are correlated with early increased CSF Aβ42 levels and working memory performance following chronic treatment. Collectively, these results suggest that long-term systemic administration of MSC-EVs in aging rhesus monkeys reduces or perhaps reverses age-related cognitive decline potentially through enhanced clearance of neurodegenerative proteins and preservation of white matter integrity and functional network connectivity.

### 4.2. Long-Term MSC-EV Treatment Improves Working Memory Span and Reduces Perseveration

A decline in performance on tasks measuring working memory is a well-characterized aspect of both human (Jenkins et al., 2000; Salthouse, 1995, reviewed in Kirova et al., 2015) and non-human primate normative aging (Baxter et al., 2023; Moss et al., 2007). These declines can be observed as early as the fifth decade of life in humans and as early as 18 years of age in the rhesus monkey (Chang et al., 2022; He et al., 2023; Ibanez et al., 2019; Makris et al., 2007; Moore et al., 2006; Moss et al., 2007). The earliest changes in cognition in both humans and primates are associated with age-related dysfunction of the prefrontal cortex (Albert et al., 2007; Blacker et al., 2007; Chang et al., 2022; Dumitriu et al., 2010; Glisky, 2007; Herndon et al., 1997; Lai et al., 1995; Moore et al., 2005; Young et al., 2014). This dysfunction, which is primarily driven by alterations in white matter, manifests as impairments on tasks measuring executive function and working memory (Moore et al., 2003, 2006; Peters et al., 1994; Peters and Sethares, 2002). Here we have shown that long-term MSC-EV treatment can shift this age-related trajectory of working memory impairment in middle-aged monkeys. Treated monkeys displayed significantly improved performance on the DRSTsp (a spatial working memory task) between baseline and retesting as compared to control (vehicle) monkeys. Specifically, treated monkeys showed longer memory spans and less perseverative errors at re-testing than at baseline, while untreated monkeys had shorter memory spans and more perseverative errors. These results indicate that MSC-EV treatment may not only halt age-related declines in working memory but, in this cohort of monkeys, improve memory performance in middle aged monkeys.

A particularly interesting finding in this study was that treated monkeys demonstrated significantly decreased perseveration. Perseveration is a response pattern commonly observed in aging and is characterized by repeating a previously rewarded response strategy for additional trials even when this strategy is not effective nor rewarded (Daigneault et al., 1992). A large body of research has indicated a link between age-related changes in prefrontal areas and an increased tendency for perseverative responses during normal aging (Arnsten, 2009; Barcelo and Knight, 2002; Fristoe et al., 1997; Haaland et al., 1987; Head et al., 2009; Manes et al., 2002; Moore et al., 2023; Raz et al., 1998; Stuss et al., 2000). This same inefficiency in cognitive shifting has also been observed in middle-aged and aged monkeys performing the DRSTsp and following lesions of prefrontal cortices (Moore et al., 2009; Moss et al., 1997). Our previous studies also demonstrated that increases in perseveration, and therefore the inability to shift cognitive strategies, are strongly positively correlated with age beginning in rhesus monkey middle age (Moore et al., 2006; Moore et al., 2023). In the present study, our results indicate that long-term MSC-EV treatment significantly reduces the overall frequency of perseverative responses between baseline and retesting. Additionally, treated monkeys showed a marked decrease in the number of perseverative errors compared to control monkeys during testing after 18 months of treatment. While previous studies have reported increases in perseveration with aging (Fristoe et al., 1997; Gunning-Dixon and Raz, 2003; Herndon et al., 1997; Kray and Lindenberger, 2000; Moss et al., 2007; Rhodes, 2004; Ridderinkhof et al., 2002), our data demonstrate that this decline can be mitigated with a systemically administered intervention such as MSC-EVs. Similarly, while other studies have shown the potential of MSC-EVs to reduce memory impairments in rodent models of AD (Chen et al., 2021; Cone et al., 2021), our study demonstrates their efficacy as a long-term intervention against normal age-related decline in a higher order species.

### 4.3. Long-Term MSC-EV Treatment Potentially Enhances CSF Clearance of Aβ Proteins Associated with Improved Memory Performance

Aging and age-related neurodegenerative diseases are associated with accumulation of protein biomarkers of axonal degeneration, such as NFL and MBP, and neurodegenerative Aβ proteins; CSF levels of these proteins have been studied as minimally invasive diagnostic markers of neurodegeneration (Backstrom et al., 2020; Bouwman et al., 2022; Li et al., 2022; Meeker et al., 2022; Mila-Aloma et al., 2020; Santaella et al., 2020; van Engelen et al., 1992). Interestingly, in the current study, monkeys treated with MSC-EVs had significantly increased CSF levels of the Aβ42 isoform at 12 months of treatment when compared to their baseline values versus control monkeys, but no differences in CSF Aβ40, NFL, or MBP were found. Aβ40 and Aβ42 isoforms aggregate as amyloid plaques in the aged brain and are a major pathological hallmark of AD (reviewed in Gu and Guo, 2013; Ma et al., 2022). Aβ42, in particular, is the predominant, more neurotoxic isoform found in amyloid plaque depositions in AD brains while Aβ40 is only detected in a subset of plaques (Gravina et al., 1995; Iwatsubo et al., 1995; Iwatsubo et al., 1994; Mak et al., 1994; Yan and Wang, 2006). However, it remains an open question whether Aβ42 and Aβ40 CSF levels correlate with disease severity and cognitive impairment in both AD and normal aging (Ritchie et al., 2014; Southwick et al., 1996). In a study by Sturchio et al., 2021, with human subjects with brain amyloidosis confirmed by positron emission tomography (PET) imaging, it was found that higher levels of soluble Aβ42 were observed in the CSF of cognitively normal individuals than in those with mild cognitive impairment and AD. Further, the higher levels of CSF Aβ42 were associated with better neuropsychological function and increased hippocampal volume in this cohort (Sturchio et al., 2021). Additionally, studies have shown that CSF Aβ levels correlate with cognitive decline in individuals found to be of greater genetic risk for AD; decreased levels of the Aβ 42/40 ratio in the CSF were correlated with a progressive decline in memory function and high CSF Aβ42 levels predicted normal cognition in amyloid-positive individuals (Sturchio et al., 2022; Tomassen et al., 2022).

It also remains an open question whether increased levels of CSF Aβ reflect actual increased Aβ accumulation in the brain parenchyma or enhanced Aβ clearance. In human AD patients, PET-based imaging of CSF clearance and Aβ showed lower levels of brain Aβ were correlated with greater CSF clearance (Li et al., 2022). Further, PET imaging and CSF Aβ levels showed that Aβ42 progressively declined 15 and 25 years before the onset of cognitive decline (Bateman et al., 2012). A longitudinal study in cognitively normal adults (Sutphen et al., 2015) also demonstrated that reductions of CSF Aβ42 began earlier in middle age before detection of PET amyloid positivity. Accordingly, amyloid brain accumulation was detected in later middle age when CSF Aβ42 levels were low, thus suggesting that the accumulation of Aβ42 in CSF precedes accumulation amyloid in the brain parenchyma detectable by PET. Therefore, these data collectively support the idea that effective clearance of amyloid from the parenchyma and into the CSF may be important for preventing pathology and cognitive impairments associated with AD as well as normal aging.

In the current study, we found that in monkeys after 12 months of systemic treatment with MSC-EVs, CSF Aβ42 and Aβ40 levels at 12 months were positively correlated with difference scores on the DRSTsp. These results indicate that treated monkeys with higher levels of CSF at 12 months had a greater improvement in working memory from baseline to retesting at 18 months – paralleling a human study in adults with subjective cognitive impairment profiling steeper rates of cognitive decline in relation to decreased CSF amyloid levels (Verberk et al., 2020). Further, our data is consistent with the asynchronous nature of these processes shown in humans, with enhanced clearance of pathological proteins likely proceeding and driving downstream changes in brain integrity (Sutphen et al., 2015).

Similarly to humans, aged monkeys exhibit cortical Aβ plaques composed of both Aβ40 and Aβ42 isoforms (Gearing et al., 1996; Li, Z.H. et al., 2020; Norvin et al., 2015; Paspalas et al., 2018) and demonstrate age-related decreases in CSF Aβ that have been shown to correlate with impaired cognitive function (Darusman et al., 2014). However, unlike humans, rhesus monkeys do not experience the widespread neuronal loss that characterizes AD pathology (Peters et al., 1994; reviewed in Luebke et al., 2010; Peters and Kemper, 2012; Peters et al., 1996), making them a valuable model for studying age-related cognitive decline in absence of the confounding effects of extensive neurodegeneration. This preservation of neuronal architecture allows for investigation of progressive cognitive and white matter changes potentially driven by Aβ accumulation, thus providing insight into the early stages of neurodegenerative processes and cognitive decline.

While our current data point to MSC-EV mediated enhancement of Aβ42 CSF clearance as a potential neuroprotective mechanism, validation of Aβ42 and Aβ40 in brain tissue will be important for future studies. Nevertheless, it has been shown that MSC-EVs can act on microglia, pericytes, and astrocytes that mediate CSF clearance (Go et al., 2020; Lu et al., 2019; Qiu et al., 2022). Our previous work using a rhesus monkey model of acute cortical injury showed that MSC-EV treatment enhanced functional recovery after injury (Moore et al., 2019). Compared to vehicle controls, the perilesional cortex from these MSC-EV treated monkeys had increased microglial ramifications and microglial phenotypes associated with damage-clearing and anti-inflammatory functions (Go et al., 2020; Zhou et al., 2023). Specifically, MSC-EV treatment was associated with increased hypertrophic microglia expressing the complement receptor protein, C1q (Zhou et al., 2023); this microglia subtype has been shown to have a protective role for clearing pathological debris in mouse models of AD (Benoit et al., 2013). Whether MSC-EVs influence CSF levels of neurodegenerative proteins through similar microglial clearance mechanisms remains unknown. This remains an important question to address in future work. As the resident macrophage of the central nervous system, microglia are primarily responsible for clearance of waste, such as amyloid debris (reviewed in Lee and Landreth, 2010). However microglia can become dysfunctional with age, which can lead to impairments in debris clearance and promote pathological protein accumulation (Thomas et al., 2022; reviewed in Antignano et al., 2023; Neumann et al., 2009; Podleseny-Drabiniok et al., 2020). Recent transcriptomic sequencing of aged human postmortem brain tissue has also revealed microglia populations with dysregulated genes that control phagocytosis and pro-inflammatory responses (Marschallinger et al., 2020). Similarly, our studies in aged rhesus monkeys have shown that markers of microglial pro-inflammatory activation and phagocytosis increase with age and correlate with cognitive decline (Shobin et al., 2017). Future work assessing the links between Aβ accumulation, microglia activity, and MSC-EV treatment in aged monkeys will be important to decipher the potential mechanism of how MSC-EVs may enhance clearance of pathological proteins.

### 4.4. Long-Term MSC-EV Treatment Preserves White Matter Structural Integrity Associated with Increased CSF Aβ Levels and Improved Working Memory Performance

A long-standing series of our group’s studies in rhesus monkeys have shown that age-related cognitive decline is correlated with increased WM and myelin pathology (Bowley et al., 2010; Peters et al., 2000; Peters and Sethares, 2002; reviewed in Peters et al., 1996). In the current study we found that compared to vehicle control monkeys, EV-treated monkeys showed higher WM generalized fractional anisotropy (GFA) in the prefrontal and temporal regions, a measure that is highly analogous to the fractional anisotropy measure from diffusion tensor imaging (DTI) reconstruction. In human aging studies focused on healthy middle-aged to aged adults, GFA has been shown to exhibit strong age-related decline most strongly in association with callosal WM fibers within the prefrontal cortex (Tseng et al., 2021). Further, our current study shows that MSC-EV-associated preservation of WM GFA measures significantly correlate with improved performance on a working memory task, the DRSTsp, which has been shown to be impaired with age in rhesus monkeys (Killiany et al., 2013; Moss et al., 1997). The DRSTsp is primarily mediated by the prefrontal cortex area 46, which is a hot spot for age-related changes in neurons and glia in our rhesus monkey model (Moore et al., 2023; Peters et al., 1994; reviewed in Luebke et al., 2010). Therefore, the effect of MSC-EV treatment on the PFC WM GFA measures may be critical to improved performance on the DRSTsp.

Our current findings show that these GFA measures of WM integrity at 18 months were correlated with the increased CSF levels of the neurodegenerative protein Aβ42 between the 6 and 12-month timepoints. Thus, our current data suggest that MSC-EV treatment may facilitate an enhancement of CSF Aβ42 clearance at 12 months that precedes and, in turn, allows for improvements in white matter integrity, consistent with findings in humans (Sutphen et al., 2015). This notion is supported by studies with our monkey model of cortical injury that demonstrate that MSC-EVs ameliorated synapse loss (Medalla et al., 2020; Zhou et al., 2023) and myelin damage (Go et al., 2021). Although the links between myelin and Aβ pathologies remain unclear, a recent study in an AD mouse model (Depp et al., 2023) showed that myelin pathology promoted Aβ deposition by overburdening microglia with myelin debris and interfering with Aβ clearance. Thus, microglia residing within white matter tracts may play a role in MSC-EV-mediated enhancement of WM integrity. In aged rodents and monkeys, white matter microglia, which are thought to play a role in lipid metabolism and phagocytosis for removing myelin debris, are known to increase with age (Safaiyan et al., 2016; Shobin et al., 2017; reviewed in Ahn et al., 2022). Our findings suggest that MSC-EV treatment preserves WM integrity, which may be linked to the promotion of microglial clearance of pathological proteins and downstream enhancement of myelin maintenance. Thus, it will be important to histologically assess the relationships between myelin, microglia phagocytosis and Aβ accumulation in the brains of these EV treated aged monkeys.

### 4.5. Long-Term MSC-EV Treatment Effects on Large-Scale Functional Connectivity and its Relationship to Structural Connectivity

Age-related changes in WM integrity can result in altered connectivity profiles of distinct resting-state functional networks governed by multiple brain regions. Age-related increases in functional connectivity between RSNs have been reported in both normal aging human lifespan studies and populations with mild cognitive impairments (Betzel et al., 2014; Zhan et al., 2016). In our group’s previous study, we reported that weaker hippocampal to neocortical structural connections correlated with worse DRSTsp performance in middle-aged rhesus monkeys, and these structural changes were associated with increased functional connection strengths (Koo et al., 2013). Thus, age-related cognitive decline seems to be attributed to malfunction of long-distance cortico-cortical functional connections due to WM damage (He et al., 2023; Koo et al., 2013; Kubicki et al., 2019; Martin et al., 2023). Interestingly, the current study is consistent with these previous data, showing that an increase in connectivity strength between the default mode to salience network was negatively correlated with WM GFA in the right middle temporal gyrus. Indeed, MSC-EV treated monkeys exhibited greater WM GFA yet weaker default mode to salience network functional connectivity. Further, at the end of the 18-months of MSC-EV treatment, we observed weaker RSN connections between the limbic and the executive networks, covering the temporal (i.e., entorhinal cortices) and prefrontal (i.e., Areas 32, 44, and 45) cortical regions, respectively, as well as weaker RSN connections between the default mode (i.e., Area 23, ventral) and the salience (i.e., insula) networks, covering the medial and inferior temporal regions. Thus, EV treatment seems to ameliorate age-related structural-functional changes in brain networks by preserving WM integrity and concomitantly recovering between-network functional connectivity to a state more similar to young brains, which may underlie improvements in spatial working memory.

## 5. Conclusions

Overall, this study suggests that long-term systemic MSC-EV treatment in late middle-aged rhesus monkeys mitigates age-related cognitive decline possibly by enhancing the CSF clearance of neurodegenerative Aβ proteins, which correlates with enhanced white matter integrity and functional brain connectivity. Further studies at a cellular and molecular level will aim to validate and decipher the mechanisms of the effects of MSC-EV treatment shown here *in vivo*. Nevertheless, the current study underscores the potential of MSC-EVs as a therapeutic intervention for cognitive decline in aging and age-related diseases, highlighting their role in maintaining brain health and function in aging populations.

## Acknowledgments

The authors would like to thank Penny Shultz, Karen Slater, Ethan Gaston, and Rudolph Beiler, DVM, for their invaluable assistance with this project.

